# fMRI Dependent Components Analysis Reveals Dynamic Relations Between Functional Large Scale Cortical Networks

**DOI:** 10.1101/066282

**Authors:** Uri Hertz, Daniel Zoran, Yair Weiss, Amir Amedi

## Abstract

One of the major advantages of whole brain fMRI is the detection of large scale cortical networks. Dependent Components Analysis (DCA) is a novel approach designed to extract both cortical networks and their dependency structure. DCA is fundamentally different from prevalent data driven approaches, i.e. spatial ICA, in that instead of maximizing the independence of components it optimizes their dependency (in a tree graph structure, tDCA) depicting cortical areas as part of multiple cortical networks. Here tDCA was shown to reliably detect large scale functional networks in single subjects and in group analysis, by clustering non-noisy components on one branch of the tree structure. We used tDCA in three fMRI experiments in which identical auditory and visual stimuli were presented, but novelty information and task relevance were modified. tDCA components tended to include two anticorrelated networks, which were detected in two separate ICA components, or belonged in one component in seed functional connectivity. Although sensory components remained the same across experiments, other components changed as a function of the experimental conditions. These changes were either within component, where it encompassed other cortical areas, or between components, where the pattern of anticorrelated networks and their statistical dependency changed. Thus tDCA may prove to be a useful, robust tool that provides a rich description of the statistical structure underlying brain activity and its relationships to changes in experimental conditions. This tool may prove effective in detection and description of mental states, neural disorders and their dynamics.

## 1. Introduction

One of the major advantages of whole brain fMRI is the detection of large scale cortical networks encompassing different and distant brain regions. These functionally defined networks rely on temporal correlations in spontaneous and task related activity (Friston, 1994; Horwitz, 2003). Both data driven analysis and hypothesis driven analysis tools are used to characterize and detect these networks. These cortical networks and their interactions have been found to reflect different cognitive states, developmental stages and neural disorders (Greicius *et al*., 2003; Seeley *et al*., 2009; Rubia, 2012). However, although current tools are effective in identifying functional connectivity within one cortical network, these tools ignore or overlook the dependencies and interactions between these networks. The ability to detect such dependencies is especially important when examining adult brain plasticity and network-related changes induced by learning, as well as in the treatment of neural disorders.

The main data driven method used to detect functional large scale networks is independent component analysis (ICA), and specifically its application in the spatial domain (sICA), in which each voxel’s time course is treated as an example for the algorithm, and linear decomposition of temporal filters is sought (McKeown *et al*., 1998). This method has been used extensively in the detection of neural disorder related changes in resting state functional connectivity (Greicius *et al*., 2004; Calhoun, Eichele, *et al*., 2009; Krajcovicova *et al*., 2011; Gallo *et al*., 2012).In sICA, spatial independence of the cortical networks is assumed, and the algorithm is directed at maximizing this independence (McKeown *et al*., 1998; Formisano, 2002). This assumption has been challenged in the context of interpreting brain activity data (Friston, 1998; Smith *et al*., 2012), and some alternative approaches have been suggested. However, these either compromise the data driven nature of ICA by using a semi-blind approach (and not overcoming the problem) (Calhoun *et al*., 2005), or undermine the robustness of ICA by examining temporal ICA, which introduces a dimensionality problem since number of voxels is much larger than number of time points sampled in fMRI (Calhoun *et al*., 2001a).

In DCA it is assumed that the components underlying the observed dataset are *dependent* on each other. Thus DCA allows cortical areas to be part of multiple cortical networks, and provides a dependency structure between these networks. This method was initially suggested and applied to examine the statistical features underlying natural images (Zoran & Weiss, 2009) by assuming a tree graph structure describing the inter-dependencies between the components (tDCA).

Here we applied this method to fMRI data, as it allows detection of overlapped networks, while still using the spatial domain and not the temporal domain and avoids the dimensionality problem. Moreover, this method uncovers not only large scale networks, but also their statistical relations and dependencies; i.e., information that is missing completely from other functional connectivity approaches. We used DCA to detect changes in cortical networks and their relations in three audiovisual experiments. In all the experiments, the auditory and visual stimuli were kept intact, but their novelty, information and task relevance were modified. This was enabled by using a visual to auditory sensory-substitution-algorithm (SSA), which is usually employed as a rehabilitation tool for the blind (Meijer, 1992). In SSA, visual information is captured and transformed to auditory soundscapes according to a set of principles. Subjects were scanned in a passive paradigm before learning SSA, in the same passive paradigm after learning SSA, and while performing an audiovisual integration task. DCA was thus used to examine how the changes in experimental context affect functional networks and their interactions.

## 2. Methods

### 2.1. tDCA algorithm

tDCA takes a similar approach as spatial ICA in assuming that the data, i.e. the voxel’s time course of fluctuations in the BOLD (blood oxygen level dependent) signal, is generated from a linear mix of basis function (which can be referred to as “causes” or “sources”). The mixing coefficients reflect how strongly each source is manifested in a specific voxel’s time course. These can be depicted as a cortical parameter map showing which cortical regions are associated with a specific source. In sICA these components are assumed to be statistically *independent*, and the spatial overlap between components is minimized. By contrast, in tDCA, components are assumed to be statistically *dependent*, in a tree dependency structure. The algorithm’s objective is to determine these basis functions or sources (or their inverses, the filters) along with the distribution of mixing coefficients and their dependency structure. We followed the methodological steps proposed by Zoran and Weiss (Zoran & Weiss, 2009) used in the context of natural images. Their method includes a preprocessing step of whitening the data, followed by an iterative process of learning the set of linear filters, the density model for the pair-wise statistics and the dependency structure itself. These steps are detailed below.

The first step is whitening the data. This is a common preprocessing operation which discards all second order correlations in the data. This allows us to focus on higher-order dependencies which are the outcome of the non-Gaussian structure of fMRI signals (Dinov *et al*., 2005). The whitening process is described briefly below.

Let the time courses of all voxels be arranged in columns to form a matrix **X**. The first step is to whiten matrix **X** such that:

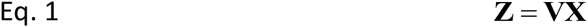

where **V** is:

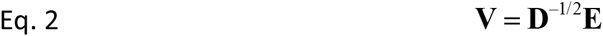

and where **E** is the eigenvector matrix of the covariance of **X** (PCA matrix), and **D** is a diagonal matrix with the corresponding eigenvalues on its diagonal. In this stage we also reduce the dimension of **X**, using only a subset of the PCA components.

Next, in order to learn the complete model we need to learn a set of linear filters, the density model for the pair-wise statistics, and the dependency structure itself. All three are learned in iterative manner. Starting with a random initialization of the parameters we repeat the following steps in iterative manner.

Learning the filter matrix: Given whitened, dimensionally reduced time courses **z**, the current estimate for the tree structure T and the pairwise density model *p*(*y_i_*, *y_j_*) (both described below), we learn an orthonormal matrix **W** such that:

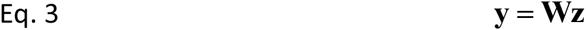

where y is the resulting coefficient vector. The distribution of the components of **y** is described using a tree-dependent model of the form:

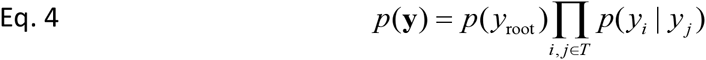

where root denotes the root node, *i, j* in *T* are the node pairs that are connected in the learned tree structure *T*. W is learned by performing a gradient ascent on the log likelihood of the model over all patches in the dataset – see Zoran and Weiss for more details.

The resulting matrix W can then be used to extract response maps by correlating its columns with the dataset, as is normally done in ICA.

Learning the tree structure: after updating the filter matrix W we learn the optimal tree structure using the Chow-Liu algorithm by relying on the pairwise mutual information (MI) between the mixing coefficient matrix (Chow *et al*., 1968). This guarantees that we learn the optimal tree with respect to the current estimate of the parameters.

Learning the pairwise density model: since we restrict the dependency structure to trees, we are faced with the challenge of handling nodes that are connected by the tree, but are in fact independent, or less dependent on one another. In order to allow for this, our pairwise density model comprises a linear mix of a factorial distribution (i.e., the coefficients are independent) and a radial non-Gaussian distribution (the coefficient are maximally dependent in our model). The mixing parameter *β* is learned for each of the edges in the tree independently.

### 2.2 fMRI Experiment

#### 2.2.1 Experimental design

Subjects were scanned in fMRI before and after learning SSA (1 hour learning session) in three experimental conditions, in order to examine the dynamic and context-dependent nature of the audiovisual integration system. All experiments included blocks of auditory stimuli and blocks of visual stimuli, delivered with different presentation rates (e.g. auditory blocks repetitions and visual blocks repetitions were not the same), in a semi-overlapped manner (Hertz & Amedi, 2010). In the first experiment (‘Pre’) blocks of visual images repeated 21 times while auditory blocks repeated 14 times. Visual blocks included three shapes (a circle, a horizontal line and a staircase); each was repeated 4 times, totaling 12 seconds per block. Auditory blocks included 3 soundscapes, which were the SSA translation of a circle, a horizontal line and a staircase. Each soundscape repeated 4 times, totaling 12 seconds per block. The rest between auditory blocks was 15 seconds, and was 6 seconds between visual blocks. In this experiment subjects were instructed to maintain fixation on a red cross in the middle of the screen and passively attend to the stimuli.

After learning, an active (‘Plus’) experiment was carried out. In this experiment auditory soundscape blocks and visual image blocks were presented to the subjects, auditory blocks repeated 20 times and visual blocks repeated 15 times. Auditory blocks included 4 soundscapes, each repeated 3 times, totaling 12 seconds per auditory block, which was followed by 6 seconds of rest. Visual blocks included 6 images, each repeated 3 times, totaling 18 seconds per visual block, which was followed by 6 seconds of rest. Subjects were instructed to press a button when they perceived a combination of a vertical line and a horizontal line, one via a visual image and the other via an auditory soundscape that together formed a multisensory “plus” (+) sign. These “plus” events occurred 10 times during this experiment as shown in the green rectangle in Figure 1D.

**Figure 1.**
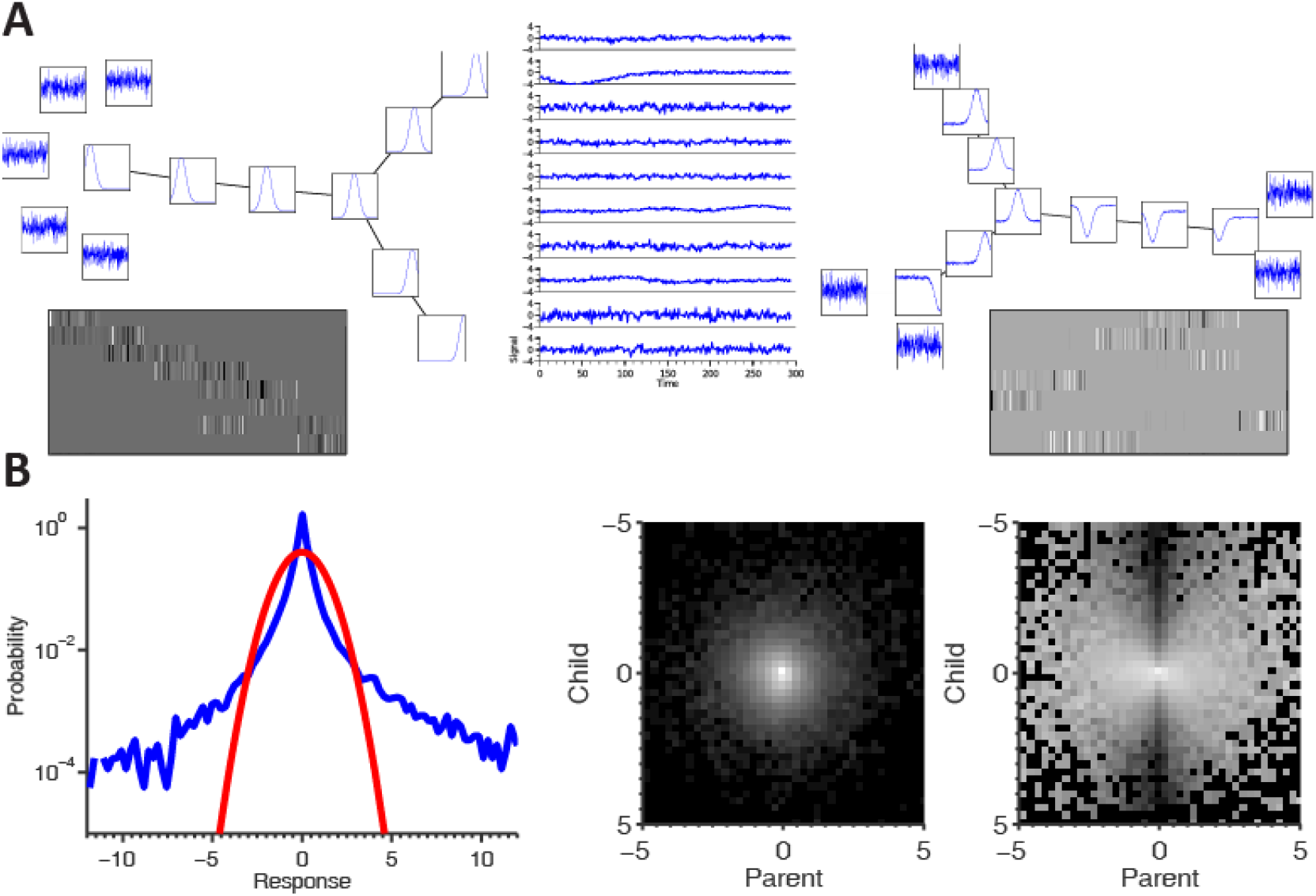
– Tree dependent component analysis. (A) Simulation results. Left: the generating tree, including basis functions and noise components. Black edges in the tree correspond to β = 1 and white edges (which are invisible) correspond to β = 0, grays are in-between values. Below it are the response maps in the brain generating the “time courses” for the simulation. Middle: some of the samples generated. Right: the learned tree model and components. Note that both the tree structure and components (including the separation from noise) were correctly learned. Below are the response maps of the non-noise components where each row of the image represents a response map to one component. Note that the ordering (and sign) is arbitrary, but that the resulting response maps indeed correspond to the “real” ones. (B) The density model used for describing the bivariate statistics of the model. Left: marginal histogram (log scale) of the 10^th^ PCA component from one of the datasets used. Note the heavy tail compared to the depicted Gaussian. Middle and Right: joint and conditional histograms of a pair of ICA components from the dataset - note the “bow-tie” shape in the conditional histogram, showing the kind of dependence we need to capture with the density model. Also note that these are ICA component responses that illustrate the failure of the ICA to remove all dependencies from the data.

The final experiment was a repetition of the ‘Pre-Passive’ experiment described above, after learning SSA (‘Post’). The same visual and auditory blocks as in the ‘Pre’ experiment were delivered, and no active response was required. Subjects were instructed to maintain fixation on a red cross in the middle of the screen and passively attend to the stimuli.

The SSA used in this study was ‘the vOICe’ developed by Peter Meijer (Meijer, 1992). This is a sensory substitution algorithm designed to provide visual information to the blind via auditory input. Time and stereo panning constitute the horizontal axis in the sound representation of an image, tone frequency makes up the vertical axis, and loudness corresponds to pixel brightness. It is hard to decipher visual information from soundscapes (SSA translation of images to sounds) before learning the principles of SSA, but even a short learning period is enough to extract visual information and detect simple objects (Kim & Zatorre, 2008; Striem-Amit *et al*., 2011).

#### 2.2.2 Subjects

A total of 11 healthy subjects (5 males, 6 females) aged 22–30 with no neurological deficits were scanned in the current study. The Tel–Aviv Sourasky Medical Center Ethics Committee approved the experimental procedures and written informed consent was obtained from each subject. We had to reject the data from one subject’s post learning passive experiment because of a technical failure of the scanner’s auditory system.

#### 2.2.3 Functional and anatomical MRI acquisition

The blood-oxygen-level-dependent (BOLD) BOLD fMRI measurements were conducted with a GE 3-T echo planar imaging system. All images were acquired using a standard quadrature head coil. The scanning session included anatomical and functional imaging. 3D anatomical volumes were collected using a T1 SPGR sequence. Functional data were obtained under the following timing parameters: TR = 1.5 s, TE = 30 ms, FA = 70°, imaging matrix = 64×64, FOV = 20×20 cm. Twenty-nine slices with slice thickness = 4 mm and no gap were oriented in the axial position, for complete coverage of the whole cortex and scanned in an interleaved order. Each experiment had 254 data points. The first eight images (during the first baseline rest condition) were excluded from the analysis because of non-steady state magnetization. Functional and anatomical datasets were normalized and aligned to standard Talairach space. Cortical reconstruction of anatomical data included the segmentation of the white matter by using a grow-region function embedded in the Brain Voyager QX software package.

### 2.3 tDCA of fMRI data

#### 2.3.1 single subjects

tDCA and ICA were carried on single subjects’ data. The BOLD signal over time (time course) from each voxel was used as a sample for tDCA analysis, with ~100000 time courses per subject. All scanned voxels were used, even if they were sampled outside of the brain, as they help estimate non-neuronal noise (see Results). Preprocessing steps included head motion correction, linear trend removal, slice time correction and high pass filtering (> 4 cycles per experiment), and spatially smoothed (spatial Gaussian smoothing, FWHM = 4mm). Functional data were spatially normalized and aligned to standard Talairach space. All these steps were carried out using the BrainVoyager QX software package. Further analysis steps were carried out using code developed in the lab in Matlab (MathWorks, Natick,MA). These steps included whitening of the data using PCA and a dimensionality reduction step in which only the first 30 PCA components were kept.

tDCA was carried out on the whitened and dimensionality reduced dataset as described above, yielding 30 filters along with their dependency tree. Each filter was used to construct a response map. The time course was used as in a standard GLM analysis (Friston *et al*., 1994), and was calculated on the original dataset. Note that the sign of the filters is arbitrary because the density model is symmetric around 0. Furthermore, negative responses should not be confused with deactivation or negative BOLD responses since they describe the relation to the corresponding component and were not compared to baseline activity (as in GLM, for example).

Two indices were devised to quantitatively estimate the temporal characteristics of the components and detect noisy components (McKeown, 2003). The first was a spike index, detecting whether a component’s time course contains a spike in time:

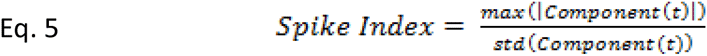

The higher this index, the higher the spike the time course contains, and the less likely it derives from neuronal activity. Spikes higher than 5 standard deviations from the baseline activity are related to head movement or scanner noise.

The second index quantified the relations between the high and low frequency energies of a component’s time course. The functional BOLD signal is characterized by high energy in low frequencies (<0.1Hz), whereas physiological noise is characterized by high frequencies (McKeown, 2003). The energy index measures this relation:

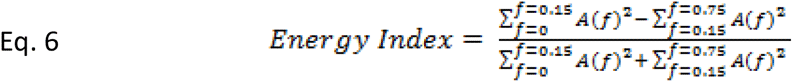

where A(f) stands for the Fourier coefficient in frequency f. When the energy index is negative, most of the energy is concentrated in high frequencies, and the time course is less likely to represent neural activity. The higher this index from zero, the more low frequencies dominate the time course spectrum, and it is more likely to derive from neural activity.

We ran ICA on single subjects’ datasets after the same preprocessing steps. The FastICA algorithm implemented in Brain Voyager software (Formisano *et al*., 2004) was used to obtain 30 components and their response maps. In order to identify similar components across the analysis results, a cross correlation matrix was calculated between all the response maps of the two sets of components. These correlation matrices were then ordered using hierarchical clustering based on the Euclidean distance between rows of the matrix, resulting in a dendrogram depicting the most similar components (using standard Matlab functions dendrogram). This enabled identification of paired components across analyses.

#### 2.3.2 Group tDCA

In order to examine the group level results we followed the concatenation method introduced by Calhun et al. (Calhoun *et al*., 2001b). In this approach tDCA is performed on a dataset created from the concatenation of all single subjects’ datasets, which results in group components and a dependency structure. Afterwards a dimensionality reduction was carried out similar to the single subject case described above. We used the first 30 PCAs from each subject, thus remaining with a whitened, dimensionally reduced time course length of 30 (this is the same as the whitened time course **z** described above). These time courses were concatenated, creating one time course length [Number of Components]*[Number of Subjects], in our case 30*11 = 330. tDCA was conducted on the concatenated dataset as was done on the single subjects. This step resulted in 30 components and their dependency structure. We dissected the group filters to get the individual filters, and reconstructed the single subjects’ response maps for each component. These were used to create a random effect measure of the components’ response maps (similar to GLM random effect analysis (Friston *et al*., 1999)). These were also the basis for comparisons of components between experimental conditions, using student t-tests between the components. Group ICA was also carried out on the concatenated dataset in the same manner. The FastICA algorithm implemented in Brain Voyager software was used to obtain 30 components and their response maps.

Group results were compared between experimental conditions and between analysis methods. As described above, the cross correlation matrix was calculated between all the response maps of the two groups. These correlation matrices were then ordered since paired components between groups were identified.

## 3. Results

### 3.1 Simulation

To validate this new method, well-controlled simulated data needs to be used to test how well it can recover the correct tree structure, and the different “activity maps” corresponding to the regions from which we created the sample. We created a simple “brain” consisting of seven different areas. Each of the areas had different characteristics, so the “time courses” generated by the voxels in this area differed. To simulate this, we created a basis of 8 time courses and 5 noise components, all connected by a tree dependency structure (because the noise is independent, it is disjoined from the tree). Each of the regions generated activations from 2000 voxels using a pair of connected nodes in the tree sampled from the density model, as well as independent Gaussian noise. We then ran the analysis steps on the simulated data, learning the tree structure, the basis functions and their corresponding component (“activity maps”), so these could be compared with the predefined underlying structure the sample was created from.

Figure 1 shows the results of the entire experiment. On the left, the generating basis functions as well as the noise components can be seen, connected by the tree. Below are the “response maps” for all components. Each row in the matrix corresponds to the activity of a single component, and each column is a different voxel. It can be seen that in different areas different components are active. In the middle, we show some actual time courses from the sample set subjected to the algorithm. On the right the learned structure can be seen. Both the tree and components were learned correctly, along with the 5 disconnected noise components. We set the gray level of each edge in this plot to be proportional to the value of beta for the edges, where the white indicates a beta very close to 0, and black very close to 1. Note that the noise components are in fact part of the learned tree, but because their *β* values (which, as noted above, measure the degree of dependency of two coefficients connected by an edge) are very close to 0, they can barely be seen. In addition, when applying the learned filters on the sample set, the same response maps are approximately recovered, with a small amount of noise.

### 3.2 tDCA of Single subject fMRI data

We began the close inspection of how tDCA performs on real fMRI dataset with single subject datasets from an audiovisual experiment. In this experiment visual images and auditory soundscapes (visual transformation to sounds via sensory substitution algorithm, see methods) were presented while the subjects lay in the scanner and were instructed to maintain fixation on a red cross in the middle of the screen. tDCA resulted in 30 components for the subjects and their dependency structure. Each component was characterized by a temporal pattern (the component’s time course) and a component response map (each voxel’s response to the component’s time course).

Two indices were used to characterize the components’ time courses based on their temporal features: an energy index and a spike index (see Methods). Figure 2A show the components of one subject in index space. Components with negative energy index have more energy in high frequencies than in low frequencies (Component 13 in figure 2A,B), and are related to physiological noise such as heart rate and respiration (McKeown, 2003). These were color coded in grey. The higher the energy index, the more low energy it contains (Component 6 in figure 2A, B), and the more likely it is to derive from neural activity. These were color coded from white (zero index) to yellow. The second index was a spike index, measuring how high a spike was above baseline. Time courses dominated by spikes (Components 12, 17 in figure 2A,B) usually derive from head movements or scanner related noise (McKeown, 2003). Components were color coded based on their spike index: yellow components had spike index of one, and faded as the index increased.

**Figure 2.**
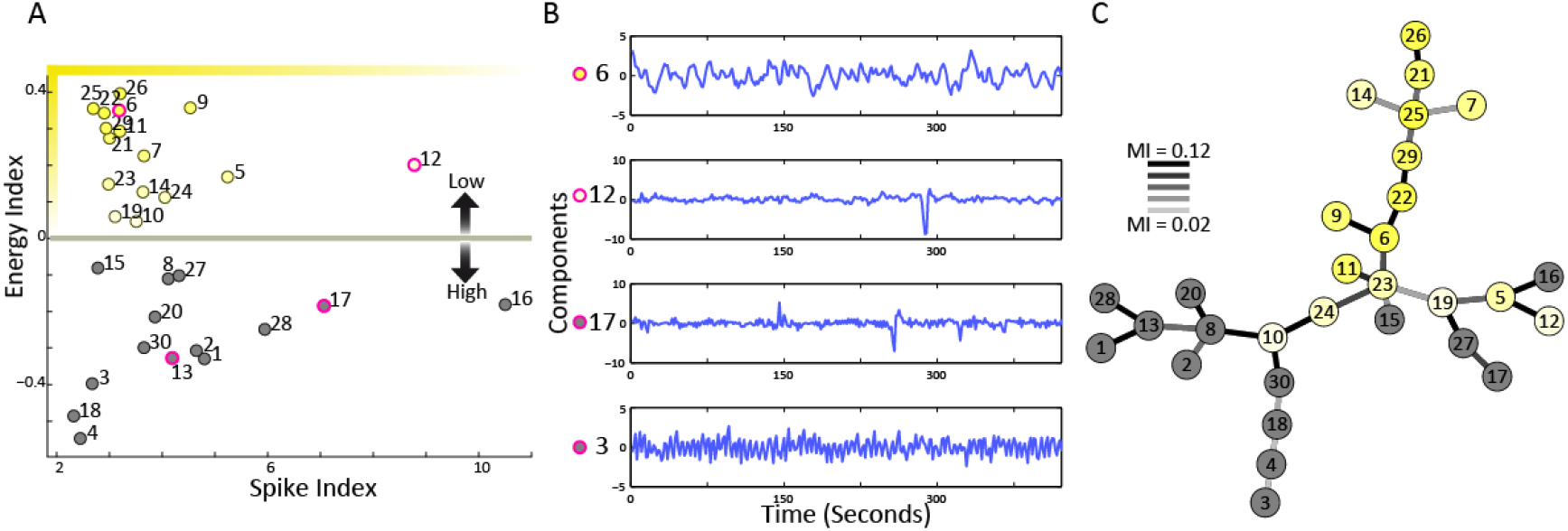
- Single subject fMRI results – temporal noise index and tree dependent dependencies. (A) The resulting DCA components of one single subject were analyzed to detect temporal noises. Two indices were assigned to each component. Theenergy index (on the y axis) is the contrast between the energy in low frequencies (f < 0.15 Hz) and in high frequencies. The spike index (on the x axis) divides the maximum value of the component with the standard deviation of the signal, and a high spike index indicates the component is dominated by one or more spikes. The circles are the detected DCA, which were color-coded based on their index values – all components with a negative energy index were colored grey, and positive energy indices were color coded on a sliding scale from white to yellow, with the brightest yellow indicating the highest energy index and lowest spike index (top left corner), and white indicating a decrease in the energy index and an increase in the spike index. (B) 4 components illustrate the different temporal characteristics associated with the temporal noise indices. These components are marked with magenta circle in the index graph in A. Component 6 had a high energy index and a low spike index, and overall low temporal noise. Components 12 and 17 had high spike indices, and presented single or several spikes in the temporal domain associated with head movements. Component 3 had a negative energy index, and demonstrated a high temporal frequency which is associated with physiological noise (breathing, heart rate). (C) The resulting tree dependent structure between components. The tree structure relies on the mutual information between the component response maps. The lines connecting the components are color coded (black to white) based on their mutual information (MI), normalized according to the maximum MI value detected (here maximum MI was 0.12). The components are colored based on their temporal noise indices, as shown in A. Interestingly, the temporally non-noisy components are clustered together on a branch of the tree, with lower indices towards the fringes of this branch, even though tree structure was determined by the spatial relation between component response maps. This clustering was evident in all the datasets examined in this study (see 10 more single subjects results in **Supplementary Figure 1**, and group results in **Figure 4**).

The nodes of the tree structure learned using tDCA (each represents a component) were colored based on the temporal indices as described above (single subjects in figure 2C, and 11 subjects in Supplementary Figure 1). Remarkably, all the yellow components, which are likely to derive from neural activity, were clustered together on a subset of the tree, and the grey components (noisy) were in the fringes of the tree. Typically, a white component (low energy index or high spike index) was adjacent to the noisy components (in grey), and a core of functional components were surrounded by white components whereas the grey components were farther away. This was the case in all the tree structures examined in this study, both in single subjects and in group analysis (Supplementary Figure 1, figure 4). This is especially remarkable since the tree structure was based on the components’ response maps, and did not take into account any temporal feature of the components’ time courses. This thus strengthens the argument that the tree structure can capture a meaningful representation of detected components, in that adjacent components share temporal characteristics.

We compared the single subject tDCA components with the most spatially similar ICA components of the same subject (see Methods). Some components showed a high similarity between methods; for example the auditory component (R = 0.72, component 9 in Figure 3) and default mode network (DMN) component (R = 0.75, component 26 in Figure 3. Interestingly, both ICA and DCA detected a highly similar CSF (cerebrospinal fluid) component localized in the ventricles and sinuses. This component had temporal characteristics similar to the neural derived components, and could only be identified as noise based on its response map. Other components that most closely resembled the ICA components did not show such high similarities. For example, the Parieto-frontal tDCA most similar ICA component correlation coefficient was only 0.46 (component 21, Figure 3). One possible explanation for this variation is the fact that tDCA allows for spatial overlap between components. This subject’s Parieto-frontal component and DMN component highly overlapped, and were also adjacent in the tree structure (Figure 2C, components 21 and 26). ICA is aimed at minimizing spatial overlap, and therefore cannot provide two components to pair with the two overlapped DCA components. Allowing spatial overlap between components led to a dramatic change in the resulting components revealing a broader range of functional cortical networks.

**Figure 3.**
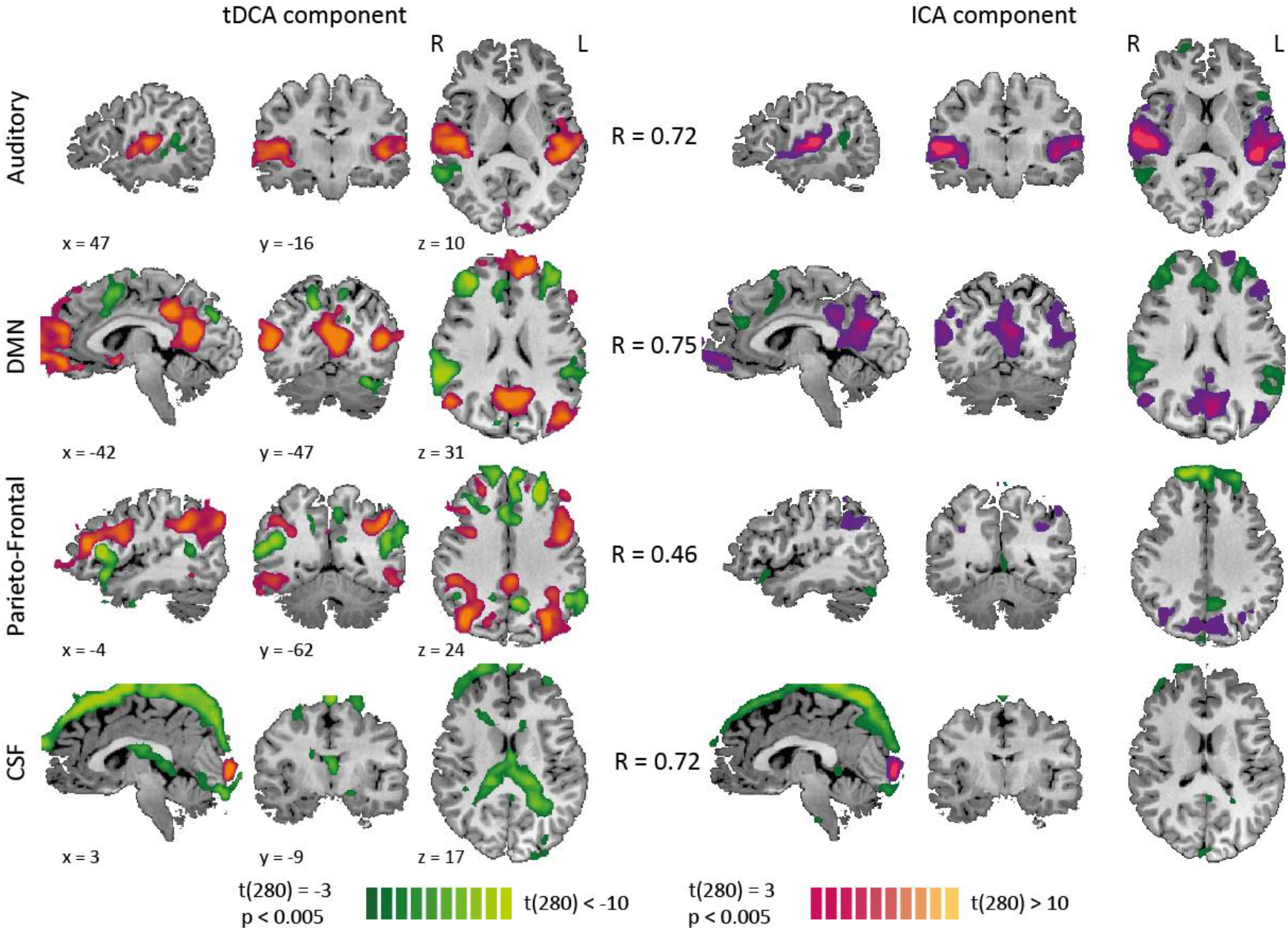
– Single Subject fMRI results – tDCA and ICA components. All component response maps were thresholded at t(280) > 3, p < 0.005). The top two component maps, 9 (auditory) and 26 (default), were highly correlated to an ICA component map (R = 0.72 and R = 0.75 respectively), and demonstrated high spatial similarities. Third component map – 21 (fronto-parietal) most similar ICA component map was not as highly correlated (R = 0.46), showing that ICA failed to detect this network. This could be explained by the fact that components 21 and 26 have high MI (0.12), and are in adjacent nodes in the tree model (Figure 2C), and the ICA algorithm is aimed at minimizing MI between components. Lowest component maps – 7 (CSF) demonstrate that even though some components did not demonstrate temporal noise, as detected by temporal noise indices, they were not derived from neural activity because they are localized in areas without neurons, in the CSF demonstrated here or in white matter. ICA also detected the CSF component (pairwise R = 0.72). CSF proximity to cortex may be the basis for its MI with non-noisy components (albeit relatively low MI).

### 3.3 tDCA of group fMRI data

Group analysis is extremely valuable in fMRI studies, because it overcomes individual subject variability and noise, and increases the signal to noise ratio. It is especially useful when trying to compare populations, for example in diagnosing clinical conditions, or in our case with the same group before and after learning novel audiovisual coupling principles. However, group analysis of ICA in general, and our approach in particular, is still a topic of debate in the literature (Calhoun, Liu, *et al*., 2009). Because components are learned independently for each subject, they vary in temporal structure and in cortical maps. This is because one subject’s cortical component map may be detected as two separate components and maps in another subject. Moreover, each subject has his/her own tree dependent structure, based on individual components. We chose to follow the approach described in Calhoun et al. (Calhoun *et al*., 2001b) for group analysis of ICA data, using concatenation of data across subjects (see Methods).

We carried out group tDCA on datasets of 11 subjects who participated in the three audiovisual experiments. These experiments were designed to examine the effect of context, e.g. task relevance, information and novelty, on audiovisual processing. This was achieved by using visual to auditory sensory-substitution-algorithm (SSA), which transforms visual images to auditory soundscapes according to a set of principles. In all three experiments auditory soundscapes and visual images were kept the same, but experimental context was modified. A ‘Pre’ condition was carried before learning SSA using a passive paradigm. A ‘Post’ condition used the same passive paradigm after learning SSA. In a third condition, ‘Plus’ condition, carried after learning SSA, subjects had to integrate auditory and visual information. They were asked to press a button when a specific combination of auditory and visual stimuli was presented (‘Integration’) (for further details see Methods). We conducted a group analysis for each experiment separately to detect common components across experimental conditions and context dependent changes.

tDCA generated components and statistical dependency structures for each of the three experiments (Figure 4). Group components were dissected to reveal the individual components’ time courses and response maps (see Methods). Average temporal noise indices (energy index and spike index) were calculated for each component based on the single subjects’ time courses. This enabled color coding of the nodes in the tree structures, similar to the way this was done in the single subject case. Here again, the algorithm placed the noisy components in the fringes of a sub-tree containing non-noisy components, in all three experiments (Figure 4). Single subjects’ response maps were evaluated statistically (random effect analysis, see Methods), and group response maps were obtained. We were able to label some of the components, e.g. auditory, visual, somatosensory, DMN and Parieto-frontal, based on the areas highlighted in their response map (described below).

**Figure 4.**
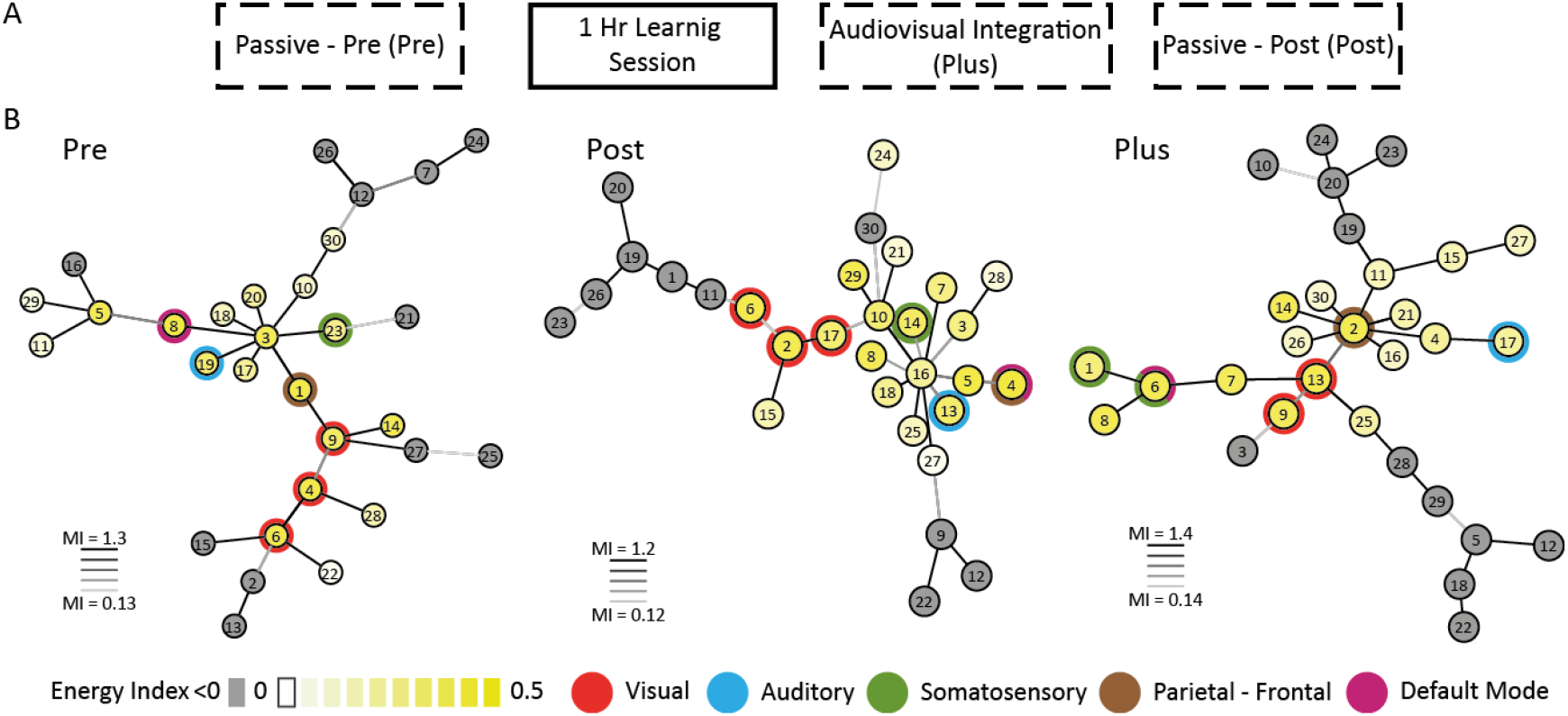
– Group fMRI results – Indices, tree structures and Positive-Negative index. (A) Order of Experiments - This study consisted of three sessions of data acquisition, interspersed by a short (1 hour) learning session outside the scanner. During the passive paradigm, visual images and auditory soundscapes were presented to the subjects in a semi-overlapped manner, while subjects were asked to maintain fixation at a cross in the middle of the screen. This paradigm was administered before learning SSA (‘Pre’), and was repeated after the learning session (‘Post’). During the audiovisual integration (‘Plus’) experiment subjects were instructed to press a button when they perceived a combination of a auditory soundscape and visual images depicting vertical line and a horizontal line, and forming a multisensory plus (+) sign. (B) Tree dependency structures of group results from three audiovisual experiments. The nodes are colored based on their average temporal node indices (see Methods), and the lines connecting the nodes are colored based on the MI between the two adjacent component maps. Here, as in the single subject case, components with less temporal noise are clustered together on a branch, even though the tree structure was determined by the spatial relations between components. Some of the nodes are also color-coded based on their corresponding visual, auditory, somatosensory, default mode and Parieto-frontal networks. In all tree structures visual nodes are adjacent.

First we compared ICA and tDCA visual components detected in the ‘Post-Learning’ experiment (Figure 5). Both ICA and tDCA detected components with response maps localized in the visual cortex. The ICA component maps had mostly positive responses, delineating one functional network, for example a component depicting primary visual area V1, from another component related to higher visual areas V2 and V3 (Figure 5A). These components tended not to overlap, as expected from ICA. tDCA components contained two anticorrelated components, with positive responses depicting one area and negative responses in other areas (Figure 5B). One tDCA component had positive responses in primary visual cortex V1 and negative responses in higher visual areas, while another had positive responses in areas with a preference for the periphery of the visual field and negative responses in areas with a preference for a foveal retinal location (Figure 5B top and middle maps) (Sereno *et al*., 1995). These results differed from the ICA components in two respects. First, networks that were detected in two different components in the ICA were detected together with the opposing response signal in tDCA. Second, the tDCA components showed high overlap, where one delineated visual cortex according to the retinal location, the other according to the processing hierarchy. This is the result of maximizing the MI between dependent components, rather than minimizing MI which is the case in ICA.

**Figure 5.**
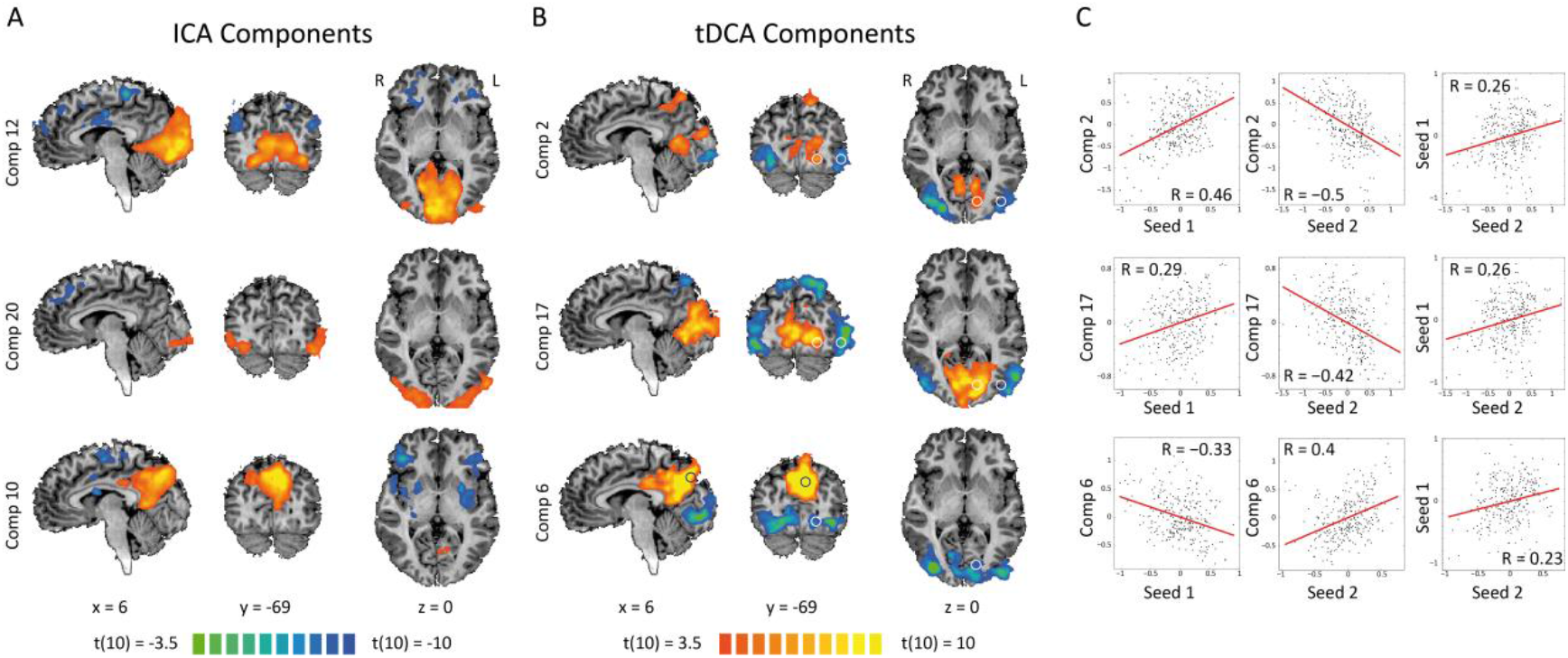
– Anticorrelations within visual system. A number of component response maps, detected in the Post-Learning experiment using tDCA and ICA included parts of the visual cortex. (A) ICA component response maps (p < 0.005). Top row depicts components that mostly include areas which prefer the periphery of the visual field, primarily in V1. Middle row depicts a component that includes a preference for the fovea. The bottom row component shows a preference for the Precuneus, outside the visual cortex. (B) tDCA component response maps (p < 0.005). Top row shows a component that elicited positive responses in periphery areas and negative responses in foveal areas. These were detected in two different ICA components. The component in the middle row elicited positive responses in V1 and periphery areas, and negative responses in higher visual areas (V3). In the bottom row is a component with negative responses in retinotopic areas and positive responses in the Precuneus. All pairs of anticorrelated responses were comprised of different dissections of the visual areas based on retinal location or processing hierarchy. (C) Time courses sampled from positive and negative responsive areas (marked with circles in B) for each component were correlated with the component time courses and with each other. In all cases the sampled time courses showed the opposite sign to the correlation with the component’s time course, but were positively correlated. The component time courses could subdivide (or spread) one functional network into two. These subdivisions were overlapped, so the visual cortex could be divided according to different features. A seed analysis would overlook these subdivisions, and ICA penalizes overlapped components.

We further examined these anticorrelated responses within one component by sampling time courses from peaks of positive and negative responses (in the circles in Figure 5B). The sampled time courses were correlated with the component’s time course and with each other. As expected, samples from positive responsive areas showed positive correlations with the component’s time course and samples from negative responsive areas showed negative correlations (Figure 5C, left and middle graphs). However, correlations between the two samples were positive (Figure 5C, right graphs). This shows that if one of the samples was used in a seed functional connectivity analysis, both areas would be part of the same functional network. tDCA components therefore managed to best spread two parts of the same network – by finding the features where two otherwise correlated areas differ the most. This is qualitatively different from ICA which finds the component best fitted a specific network. The tDCA component seems to best describe (or span) the relation between two components.

We compared components across experiments to examine which components remained the same and which experimental condition dependent changes emerged. In all three experiments, components related to sensory networks remained the same (Figure 6). The visual component, with anticorrelations between primary visual areas and higher visual areas was detected in all three experiments (p < 0.005, Figure 6 top row). Auditory components also remained constant across experiments (p < 0.005, Figure 6 bottom row). Auditory components did not demonstrate anticorrelations within the auditory cortex. Both in the auditory and visual cases, no significant differences between experiments were found when these were contrasted directly. This consistency across experiments was expected since the auditory and visual stimuli were the same (or very similar) across experiments. This consistency suggests that overall, the sensory areas were less affected by changes in experimental conditions and the context in which sensory inputs were delivered. Moreover, it demonstrates that tDCA reliably uncovered sensory components in three independent analyses of three datasets. It is important to uncover similar results using non-overlapping datasets, when establishing a novel data analysis method

**Figure 6.**
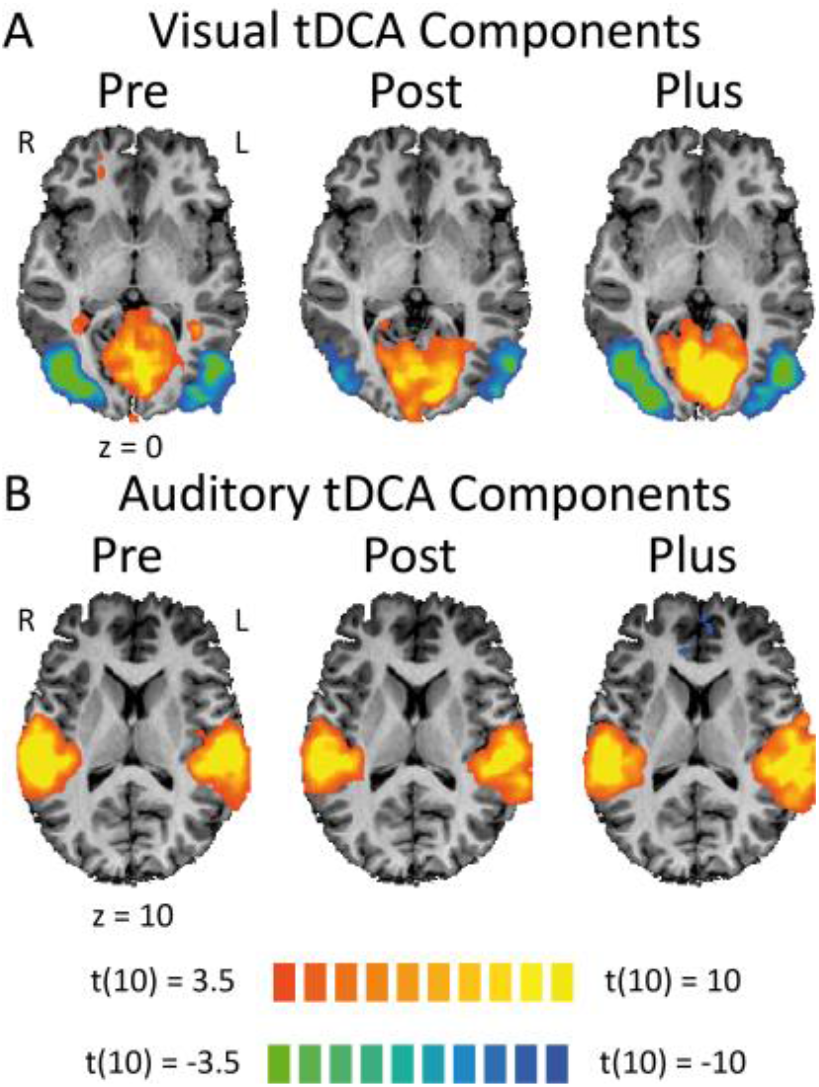
– Consistent auditory and visual components across experiments. (A) Similar visual components in all three experiments, which include anticorrelated responses in periphery and foveal areas (p < 0.005). No significant differences were detected between these maps. (B) Maps for the auditory components detected in all three experiments (p < 0.005). Here only positive responses were detected, which cover HG and PT. No significant differences were detected between these maps. Components of sensory cortices were consistently detected in all three experiments. Auditory and visual stimuli were kept the same throughout these experiments, while task, novelty and information changed.

Context dependent changes were found in the Parieto-frontal network components (Figure 7). The Parieto-frontal network is associated with control of attention and task related activity (Corbetta, 1998), and includes the bilateral Intraparietal Sulcus (IPS) and prefrontal areas (Frontal eye field FEF). The Parieto-frontal network was detected in all three experiments, including IPS and FEF (p < 0.005 Figure 7A). However, context dependent changes were detected, in that other areas became functionally connected to this network, and some areas became more or less pronounced. Direct comparison between the ‘Pre’ and ‘Post’ learning experiments revealed how the bilateral Insula became functionally connected to the Parietofrontal network after learning SSA (p < 0.005 Figure 7B left map). This could have been the result of the translation of auditory soundscape to visual framework in the Insula (Bushara *et al*., 2001), which is possible only after learning SSA principles. Comparison between ‘Post’ and ‘Plus’ conditions showed that in the absence of the task, the network was more left lateralized, whereas when the active audiovisual integration task was introduced the network was more symmetric (p < 0.005 Fuigure 7B right map). This is in line with reports that there is a more left lateralized network for object detection (Amedi *et al*., 2007), and that the right hemisphere is more involved in active control of attention (Corbetta & Shulman, 2002). All the changes in the Parieto-Frontal network were within the network; i.e., through inclusion or exclusion of areas to the network or changes in area responses, but not in its relation with other networks. Such context dependent changes can be expected in the responses of associative areas outside sensory areas, given that the experimental context changed.

**Figure 7.**
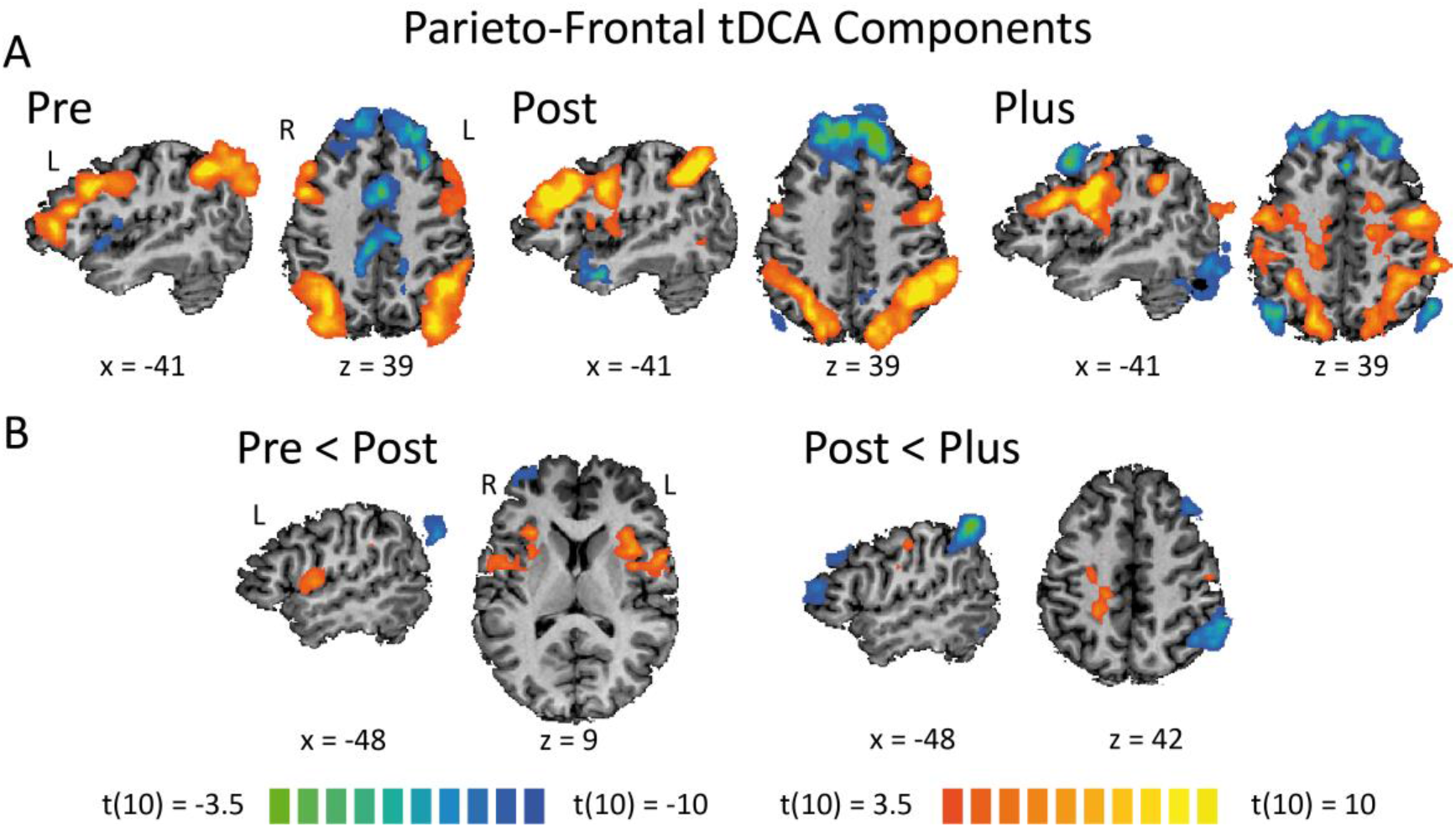
– Context dependent changes within networks: Parieto-Frontal components. (A) Parieto-frontal networks were detected in all three experiments, including IPS and FEF (p < 0.005). However these differed as a function of the changes in experimental conditions and the task, novelty and information conveyed by sensory inputs. (B) These changes are shown by contrasting components between experiments. The Parieto-frontal network was correlated with bilateral insula after learning but not before learning (left map, p < 0.005). It was also more pronounced in the left hemisphere after learning in the passive paradigm than in the active audiovisual detection experiment (right map, p < 0.005).

Finally, context dependent changes were detected in the DMN network (Figure 8). The DMN network includes the Precuneus, bilateral Temporal-Parietal Junction (TPJ) and mid-frontal cortex, and is associated with intrinsic cognitive processes, in contrast to activity related to extrinsic stimuli and task (Raichle *et al*., 2001; Goldberg *et al*., 2008). Moreover, the DMN was shown to be dissociated from a network of areas oriented toward extrinsic processes, in studies applying data-driven clustering approaches (Golland *et al*., 2008), GLM analysis (Golland *et al*., 2007) or anticorrelations (Fox *et al*., 2005). As our analysis examines relations between cortical networks the DMN and its related networks are of great interest to us. DMN components were detected in all three experiments (p < 0.005, Figure 8A – negative responses). Although the DMN network itself did not change significantly between experiments, its associated network, i.e., the network anticorrelated to the DMN in each component, changed significantly. In the ‘Pre’ condition it was anticorrelated with the bilateral Insula, in the ‘Post’ condition it was anticorrelated with Parietofrontal network which is related to attention and object detection, and in the active ‘Plus’ experiment it was anticorrelated with somatosensory areas and the insula which is related to the motor response to target detection. These changes were also observed when the components were contrasted between experimental conditions (p < 0.005, Figure 8B). These context dependent changes were not within the network detected in the component as was the case for the Parieto-frontal network, but rather were found in the relations between networks, in that the DMN was associated with different extrinsic process in each experiment. These changes followed the changes in task relevance, information and novelty of the sensory input.

**Figure 8.**
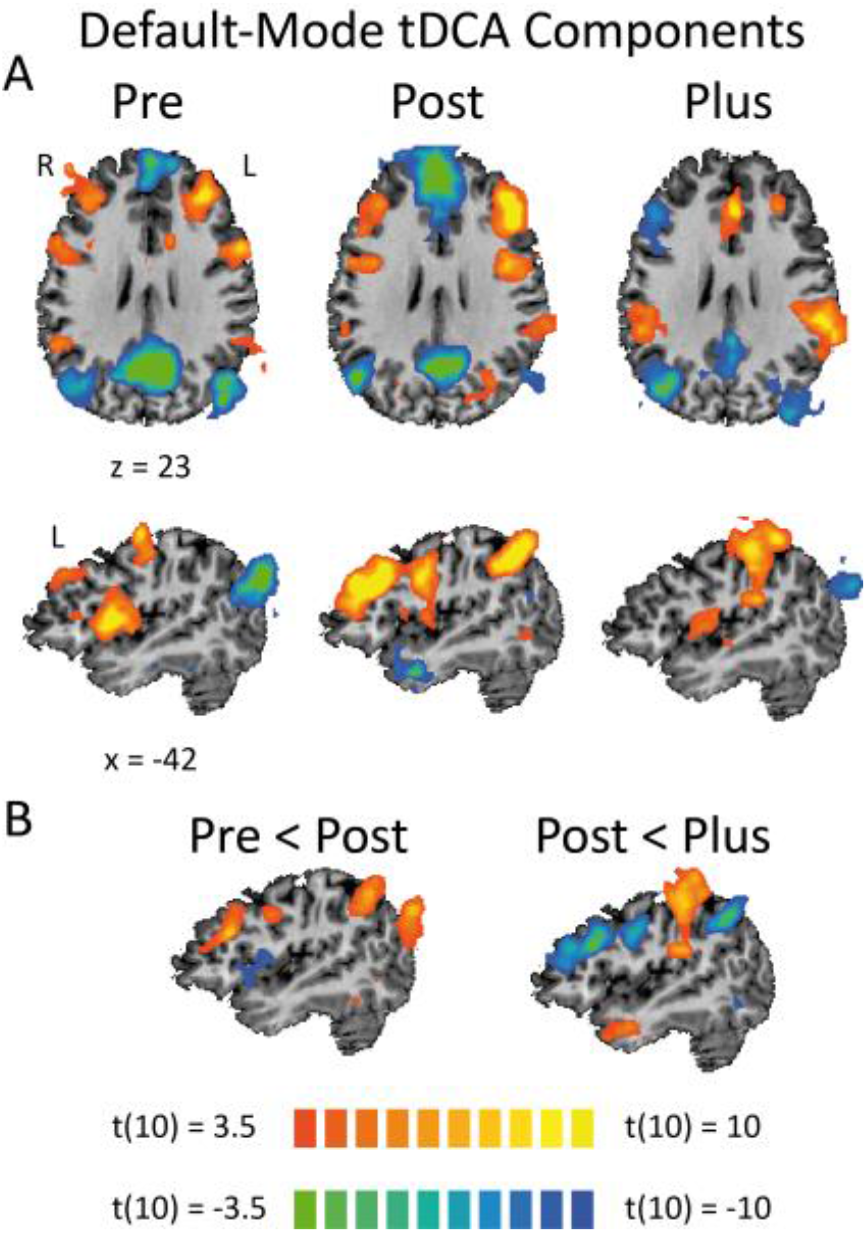
- Context dependent changes between anticorrelated networks: DMN components. (A) DMN was detected in all three experiments (p < 0.005). Whereas negative responses in DMN, including the Precuneus, TPJ and mid-frontal areas remained constant between experiments, it was anticorrelated with different networks in each experiment. Before learning, it was anticorrelated with the bilateral Insula, after learning in the passive experiment it was anticorrelated with the Parieto-frontal network and bilateral Insula, and in the active audiovisual detection experiment it was anticorrelated with left motor cortex and bilateral insula. (B) This pattern was also detected when these components were directly contrasted between experiments, showing significant changes in DMN conjugated networks, but not between DMN. This is evidence of context dependent changes in the relations between cortical networks which does not necessarily involve changes within these networks. This information is unique to tDCA.

## 4. Discussion

In this study we introduced a novel data driven approach to extract both components and their statistical dependency structure underlying fMRI data, following the algorithm introduced by Zoran et al. (Zoran & Weiss, 2009). First we demonstrated the feasibility of this method using a simulation (Figure 1), and showed that the algorithm revealed the filters and dependency structure underlying the simulated data. We then applied tDCA to study changes in cortical networks and their dependencies in audiovisual fMRI experiment datasets. We characterized temporal features of the components’ time course associated with the noisy, non-neural related signal. tDCA clustered together the non-noisy components on a subset of the tree structure, whereas the noisy components were located in the fringes of the tree, even though the tree structure was based on a spatial distribution of the components’ response maps (Figure 2, Supplementary Figure 1). When group tDCA results were compared with group ICA results, interesting characteristics of tDCA components were revealed. The first was that tDCA learns pairs of anticorrelated networks within a functionally connected network. While the ICA finds components which best characterize one area, tDCA extracts components that best spread over the two networks. This was shown in the case of visual components (Figure 5). This pair detection also results in a high overlap between components’ response maps, which was not found in the ICA, in that the algorithm tries to minimize spatial overlap between components. Finally, tDCA was used to examine how changes in the experimental context in which auditory and visual stimuli are delivered affect sensory processing in the brain. Sensory components did not change throughout the experiments because the stimuli themselves remained the same, showing that tDCA reliably detected sensory networks (Figure 6). Two types of context dependent changes were found. The first was changes within a functional network, such as the one that was found for the Parieto-frontal network, in which some areas were excluded or included in the network in different experimental conditions (Figure 7). The second type was changes in the relations between functional networks as a function of changes in the network pairs. This was seen in the case of DMN, which changed its anticorrelated networks according to the experimental condition (Figure 8). This level of information; namely, relations between functional networks, is completely absent from ICA. These results show that tDCA is a viable and meaningful data driven approach to fMRI data, and can characterize the components and dependency structure it obtains.

The use of data driven approaches to detect large scale functional networks in the brain is part of a growing interest in this phenomenon as fMRI has become an established functional brain imaging tool. In fMRI the entire brain is scanned over time, with relatively high spatial resolution. This differs considerably from electrophysiology studies that are constrained to a small area of the brain in which the electrode is inserted. This allowed for the detection of relations between distant cortical areas, and introduced the notion of functional and effective connectivity (Friston, 1994; Horwitz, 2003). Functional connectivity relates to the similarity (correlation) in temporal fluctuations in the signal between two distinct areas. The most common use of this approach is in seed analysis, in which a time course is sampled from a pre-defined cortical area, and the ‘seed’, is compared with brain activity throughout the brain. The areas which show a high correlation with the seed time course are defined as functionally connected to the seed. This approach has been especially useful in the detection of altered functional connectivity in different populations, such as during development (Rubia, 2012), in different neurological disorders (Greicius *et al*., 2004; Krajcovicova *et al*., 2011), and in different experimental conditions(Kim & Zatorre, 2011).

Spatial ICA has been suggested as a tool to extract such ‘seed’ time courses, which represent different functional networks, in a data driven manner, and hence overcome the problem of seed selection (McKeown *et al*., 1998). It has become a popular tool in detection of functional networks, especially when using a resting state paradigm (Greicius *et al*., 2004; Calhoun, Eichele, *et al*., 2009; Krajcovicova *et al*., 2011; Gallo *et al*., 2012). However, its maximization of spatial independence has been criticized as not reflecting the brain’s complexity and connectivity (Friston, 1998; Smith *et al*., 2012). Moreover, it was argued that ICA fails to maximize spatial independency (Daubechies *et al*., 2009). Temporal ICA does not make a spatial independence assumption, thus allowing for spatial overlap between components. However, since there are orders of magnitude more voxels than sampled time points in fMRI data, and ICA requires a large number of samples, this approach is not commonly used. Recently Smith et al. (Smith *et al*., 2012) were able to carry out a temporal ICA by employing an advanced fast fMRI protocol and pooling data across subjects, and successfully revealed temporal functional modes of activation. Even though sICA is more robust, alternatives are being sought which more accurately depict large scale cortical networks.

tDCA overcomes the spatial independency demands of sICA, without falling into the dimensionality problem of temporal ICA in that it uses voxels as samples, as does sICA. This means that tDCA can be used even in standard fMRI acquisition protocols, and can be carried out at the single subject level. tDCA reveals components that are characterized by pairs of conjugated, anticorrelated cortical networks. These may reflect functional connections between pairs of cortical networks. Anticorrelations between large scale cortical networks have been discussed in the context of the default mode network (DMN) and resting state fMRI (Fox *et al*., 2005). Anticorrelation was also found between a task oriented network involved primary sensory areas and attention areas, and an intrinsic network overlapping the DMN in the resting state. Other studies on this effect have reported the importance and reoccurrence of these anticorrelations as a descriptive characteristic of the relation and interactions between large scale cortical networks. For example, different anticorrelation patterns for different parts of the DMN have been identified, which suggests different roles for DMN nodes (Uddin *et al*., 2009). Another study found a disruption of anticorrelations patterns in sleep deprivation, and argued that proper interactions between large scale networks are crucial (De Havas *et al*., 2012). Although it has been claimed that anticorrelations are introduced by the effect of global signal regression (Murphy *et al*., 2009), the fact remains that some temporal components can spread two opposite signal responses in two cortical networks. Our results bear remarkably close resemblance to the temporal functional modes identified by Smith et al. (Smith *et al*., 2012) using temporal ICA (mentioned above), who found components that contain two anticorrelated networks. Conjugated pairs of large scale cortical networks may therefore reveal an important characteristic of large scale networks interactions and relations. Furthermore, the changes in conjugated pairs and dependency structure shown here provide a unique view on the dynamic nature of these interactions, in that the anticorrelations between two networks are replaced by anticorrelations with another network as experimental conditions are modified.

tDCA was used here to describe consistent and dynamic sensory processing as a function of changes in experimental conditions that affected information, novelty and task relevance of the sensory inputs. tDCA was able to uncover context dependent changes in the networks paired with a DMN component subsequent tothe changes in experimental conditions. This is a good example of the kind of information revealed by examining the relations between functional networks, which is the core idea in this analysis. Changes within functional networks, as was found in the Parieto-frontal network, could also have been found using seed analysis or ICA (however the restriction on overlapped networks can sometimes lead to failure to detect some network components). Moreover, ICA appears to be especially useful in describing one specific network, as was seen in the detection of visual components, whereas tDCA tends to find pairs of networks. It is important to acknowledge that different tools, with different assumptions and biases, lead to different descriptions of cortical activity. These should be used appropriately according to the question at hand, and are not mutually exclusive. It is important to continue to explore the statistics and features underlying fMRI data, since changing the assumptions and models may lead to novel insights, better understanding and characterization of large scale cortical activity. This may prove useful in future detection of ‘modes’ of activity, cognitive states and functional effects of neural disorders.

## Acknowledgments

This work was supported by a career development award from the International Human Frontier Science Program Organization (HFSPO), The Israel Science Foundation (grant number 1530/08), a European Union Marie Curie International Reintegration Grant (MIRG-CT-2007-205357), a James S. McDonnell Foundation scholar award (grant number 220020284), the Edmond and Lily Safra Center for Brain Sciences Vision center grant (to AA). UH and DZ wish to acknowledge the Charitable Gatsby Foundation for their support.

## Author Contributions

UH wrote the main manuscript text and prepared the figures, carried the fMRI experiment and analysis and adapted code written by DZ to fMRI data and group results. DZ carried out the simulation and prepared Figure 1. DZ wrote the original algorithm code (see Zoran and Weiss(Zoran & Weiss, 2009)). All authors reviewed the manuscript.

## Additional Information

The author(s) declare no competing financial interests.

---

**Supplementary Figure 1.**
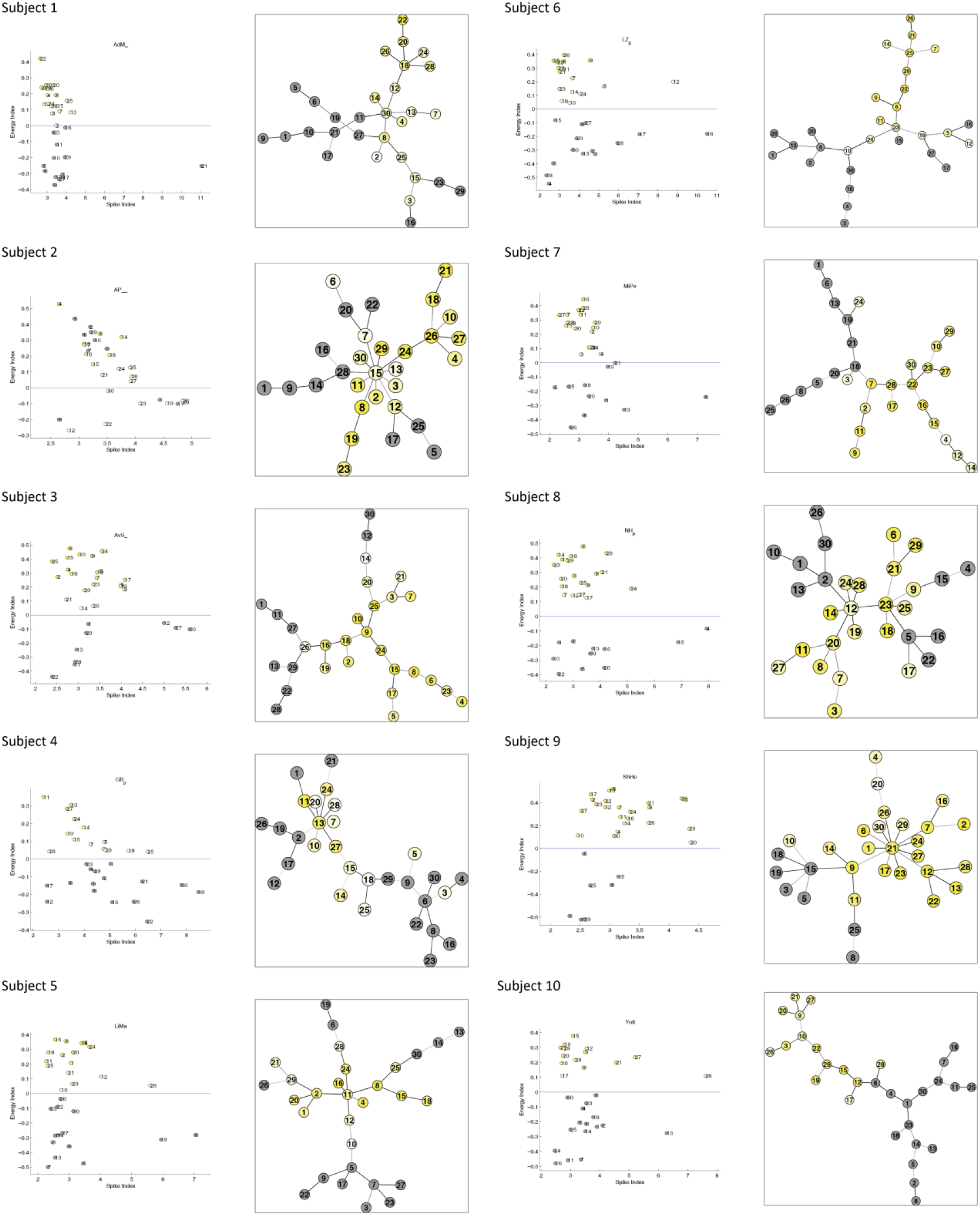
– 10 single subjects' temporal noise indices and tree dependency structures.

